# Parabrachial complex processes dura inputs through a direct trigeminal ganglion-to-parabrachial connection

**DOI:** 10.1101/2020.10.27.356972

**Authors:** Olivia Uddin, Michael Anderson, Jesse Smith, Radi Masri, Asaf Keller

## Abstract

Migraines cause significant disability and contribute heavily to healthcare costs. Irritation of the meninges’ outermost layer (the dura mater), and trigeminal ganglion activation contribute to migraine initiation. Maladaptive changes in central pain-processing regions are also important in maintaining pain. The parabrachial complex (PB) is a central region that mediates chronic pain. PB receives diverse sensory information, including a direct input from the trigeminal ganglion. We hypothesized that PB processes inputs from the dura. Using *in vivo* electrophysiology recordings from single units in anesthetized rats we identified 58 neurons in lateral PB that respond reliably and with short latency to electrical dura stimulation. After injecting tracer into PB, anatomical examination reveals retrogradely labeled cell bodies in the trigeminal ganglion. Neuroanatomical tract-tracing revealed a population of neurons in the trigeminal ganglion that innervate the dura and project directly to PB. These findings indicate that PB is strategically placed to process dura inputs, and suggest that it is directly involved in the pathogenesis of migraine headaches.

## 1. Introduction

Migraines are a highly prevalent condition involving recurrent severe headaches. These disrupt daily activities and significantly contribute to healthcare burden ^1,2^. Migraine headaches are often accompanied by autonomic instability, photophobia, phonophobia, photophobia, phonophobia and sensitivity to normally innocuous tactile or thermal stimuli (allodynia) ^3^. Allodynia impacts a significant proportion of migraine patients ^4^ and is a predictor of poor treatment response and migraine chronification ^5,6 7^.

A number of mechanisms have been identified as likely contributors to migraines (reviewed by ^8^ and^9^). A key anatomical substrate in migraine pain is the trigeminal system, that relays sensory information, including pain, from the face and head region. The outermost layer of the meninges, the dura mater, is innervated by sensory afferents of neurons in the trigeminal ganglion. Dura manipulation in humans during neurosurgery is often painful ^10^ and dura irritation is considered an initiating factor in migraine ^11^. In rodent models, dura irritation models migraine-like symptoms ^12,13^

The parabrachial complex (PB) is a bilateral midbrain structure that plays important roles in pain processing, as well as many survival-related homeostatic and interoceptive functions ^14–17^ (reviewed by ^18^). PB’s anatomical connections place it in an ideal position to modulate pain. It receives direct spinal inputs from the trigeminal nucleus and dorsal horn ^19–21^ and communicates with regions important for sensory and emotional processing ^19,22,23,24^. Amplified activity of PB neurons is causally related to chronic pain ^25–28^ and changes in PB activity contribute even more heavily in models of craniofacial and orofacial pain ^29^. Significantly, there exists a direct connection from trigeminal ganglion neurons to PB ^30–33, 29^, bypassing the canonical node in the spinal trigeminal nucleus. Thus, PB is strategically placed to process dura inputs, and to be involved in the pathophysiology of migraine.

Here, we test the hypothesis that PB processes inputs from the dura.

## 2. Methods

### 2.1 Animals

All procedures were conducted according to Animal Welfare Act regulations and Public Health Service guidelines and approved by the University of Maryland School of Medicine Animal Care and Use Committee. Twenty-two male and female Wistar rats (Envigo, Indianapolis, IN, or bred in-house), ages 3 to 12 months, contributed to the electrophysiology data presented here. For anatomical tracing studies, we used 5 Wistar rats (3 female, 2 male) ages 3 to 6 months.

### 2.2 Electrophysiology Preparation

We anesthetized (Level III-2, as defined by ^34^) all rats with 20% urethane in sterile saline. Rats were placed in a stereotaxic frame with a heating pad to maintain body temperature. We made a small craniotomy over the recording site to target PB (AP −9.2 and ML +1.9, relative to bregma, and DV −6.0 mm, relative to dura surface). We exposed the dura mater, over the middle meningeal artery, for stimulation.

### 2.3 *In vivo* Electrophysiology

Using platinum-iridium recording electrodes (2–4 MΩ) produced in our laboratory, we recorded from the PB both contralateral and ipsilateral to dura stimulation site. We isolated units responsive to electrical dural stimulation and digitized the waveforms using a Plexon acquisition system (Plexon Inc., Dallas, TX).

### 2.4 Identification of Dural Stimulus-Responsive Neurons

We used bipolar platinum-iridium electrodes to electrically stimulate the dura. We positioned these electrodes over the middle meningeal artery, either contralateral or ipsilateral to the site of PB recording. Electrical stimulation parameters were 1 to 3 mA intensity, 800 to 850 μs duration, 0.3 Hz. To define dura stimulus-responsive neurons, we used NeuroExplorer (Plexon Inc, Dallas, TX) to construct peristimulus time histograms (PSTHs), and cells that fired with greater than 95% confidence in the first 100 ms after stimulation were considered responsive to dura stimulation.

### 2.5 Histological Confirmation of Recording Site

To confirm the location of recorded neurons we made electrolytic lesions at the conclusion of recording experiments. Animals were perfused transcardially with 0.05M phosphate buffered saline, followed by 4% paraformaldehyde (PFA). Brains were extracted and incubated overnight in 4% PFA at room temperature. We then sliced brains coronally into 70 to 80 μm thick sections and stained them with Toluidine Blue.

### 2.6 Electrophysiology Data Analysis

Cells were sorted using Offline Sorter (Plexon) using dual thresholds and principal component analysis. We calculated spontaneous mean firing rates using NeuroExplorer, and used a custom MATLAB (MathWorks, Natick, MA) script to quantify responses to electrical stimulation of the dura.

We quantified as response magnitude as the probability of a PB neuron response in the 100 ms following electrical stimulation.Significant responses were defined as spikes that exceeded the 99% confidence interval of pre-stimulus, spontaneous firing rates.

### 2.7 Anatomical Tracing

We deeply anesthetized animals with isoflurane in a stereotaxic frame, and used a dental drill to expose the region over the parabrachial nucleus and over the middle meningeal artery. We used a glass pipette with a 40 μm tip to pressure inject (Nanoliter 2010 pump, World Precision Instruments, Sarasota, FL) 500 nL (50 nl/minute) of CF568-conjugated cholera toxin B (CF568 conjugated CTB; Biotium, Fremont, CA) into PB unilaterally (right side). We made a craniotomy over the right middle meningeal artery and carefully exposed the dura, applying 5 μl of Fluorogold (Hydroxystilbamine FluoroGold, Biotium, Fremont, CA) to the area, diluted to 5% in sterile saline, using a sterile pipette tip. We covered the dura application site with sterile Gelfoam and closed the incision with nylon suture. One week after tracer application we deeply anesthetized the rat with urethane and performed a transcardial perfusion with 0.05M PBS followed by 4% PFA (Sigma Millipore). We harvested the brain and trigeminal ganglia, placing them into 4% PFA overnight. We then transferred tissue to a solution of 30% sucrose in 0.05M PBS, until the tissue blocks sank. We then froze the tissue (−20°C) and used a cryostat (CM1860, Leica Biosystems, Buffalo Grove, IL) to cutsection the PB injection site and the spinal trigeminal nucleus (positive control region) at 40 μm thickness. We cutsectioned the trigeminal ganglia on a cryostat at 30 μm thickness. We then rinsed all sections in 0.05M PBS six times, before mounting the tissue on slides and cover-slipping them for imaging. We searched for and imaged labeled cells using a Confocal Microscope (SP8, Leica Biosystems, Buffalo Grove, IL).

### 2.8 Statistical Analysis

We analyzed all data using GraphPad PRISM version 8.2.0 for Macs (GraphPad Software, La Jolla, CA). When the assumptions necessary to use parametric tests were not met (normal distribution, independent data, and homogenous variance), we used non-parametric statistics. To determine age and sex differences we averaged values of each metric for an individual animal and compared these averages between sexes or ages (2.5-3 months, 5-6 months or 8-12 months).

### 2.8 Rigor and Reproducibility

We adhered to accepted standards for rigorous study design and reporting to maximize the reproducibility and translational potential of our findings as described by Landis et al. ^35^ and in ARRIVE (Animal Research: Reporting In Vivo Experiments).

## 3. Results

### 3.1 PB neurons respond to dura stimulation

We identified 58 PB neurons from 22 rats that responded to electrical stimulation of the dura mater. Responsive PB neurons were primarily located in the external lateral portion of PB (Fig. 1A). Figure 1B depicts the action potential waveforms and the response patterns of two representative neurons. The peak response latency of PB responses to dura stimulation ranged from 4 ms to 25 ms (median 10 ms, 95% C.I. 8 to 12ms), onset latency ranged from 4 ms to 20 ms (median 7 ms, 95% C.I. 7 to 8 ms), and rise time ranged from 0 ms to 11 ms (median 1 ms, 95% C.I. 1 to 2 ms) (Fig. 2A, B). Response magnitudes ranged from 0.001 to 2.1 spikes/stimulus (median 0.44 spikes/stimulus, 95% C.I. 0.26 to 0.67 spikes/stimulus) (Fig. 2D). The median spontaneous activity of these neurons was 0.72Hz (95% C.I. 0.22-3.7). Thus, PB neurons respond reliably and with short latency to dura stimulation.

**Figure 1:**
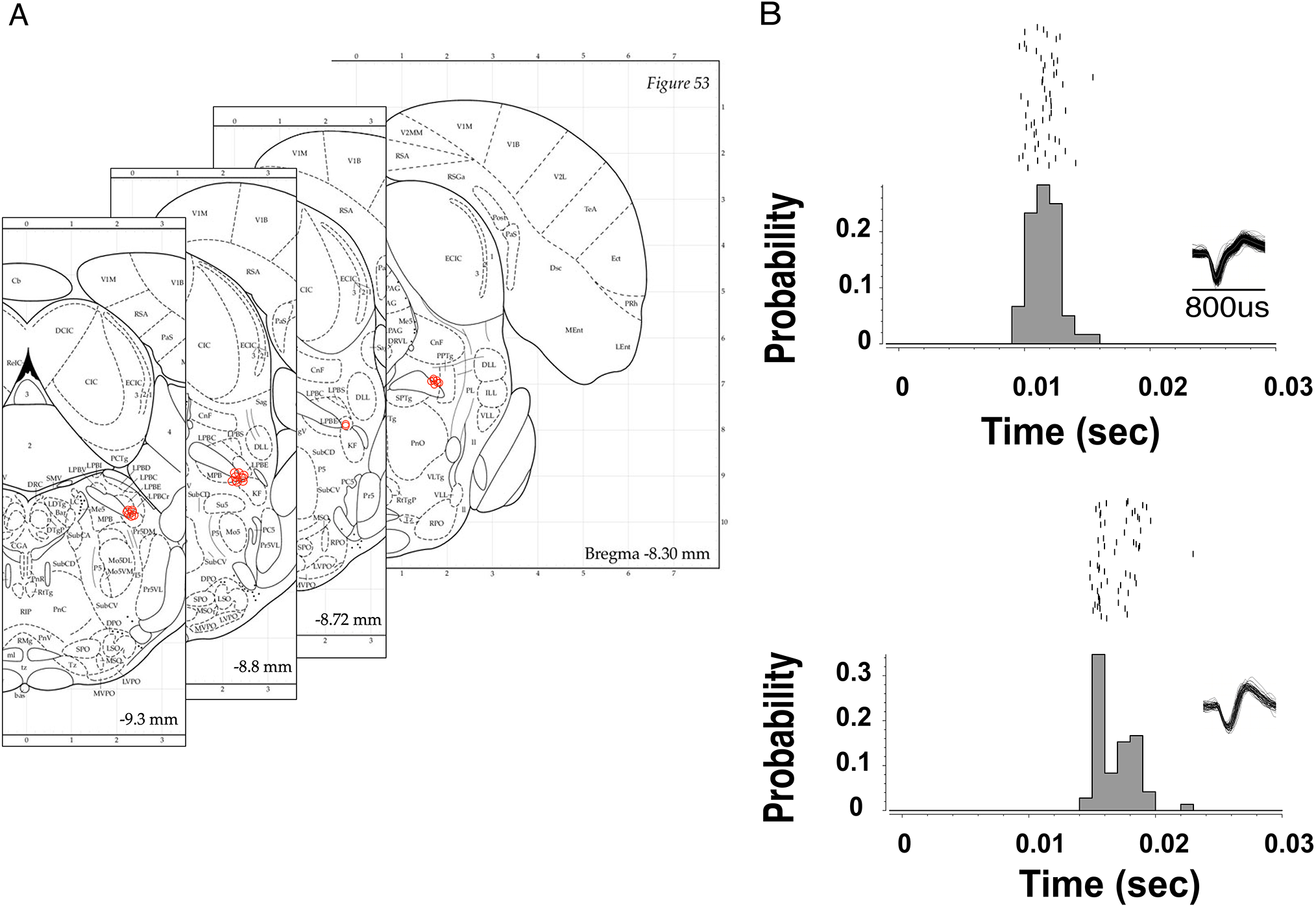
(A) Neurons responding to dura stimulation are in the external lateral PB. Atlas plates^1^ diagramming coronal sections of the rat brain, in the vicinity of PB. Red circles depict the coordinates of dura-responsive PB neurons. (B) PB neurons respond reliably to electrical dura stimulation. Two representative perievent rasters from dura-responsive PB neurons. Insets depict the waveform for each neuron. Stimulation onset is aligned to time zero. Rows of rasters above the histogram depict consecutive trials of dura stimulation.

**Figure 2:**
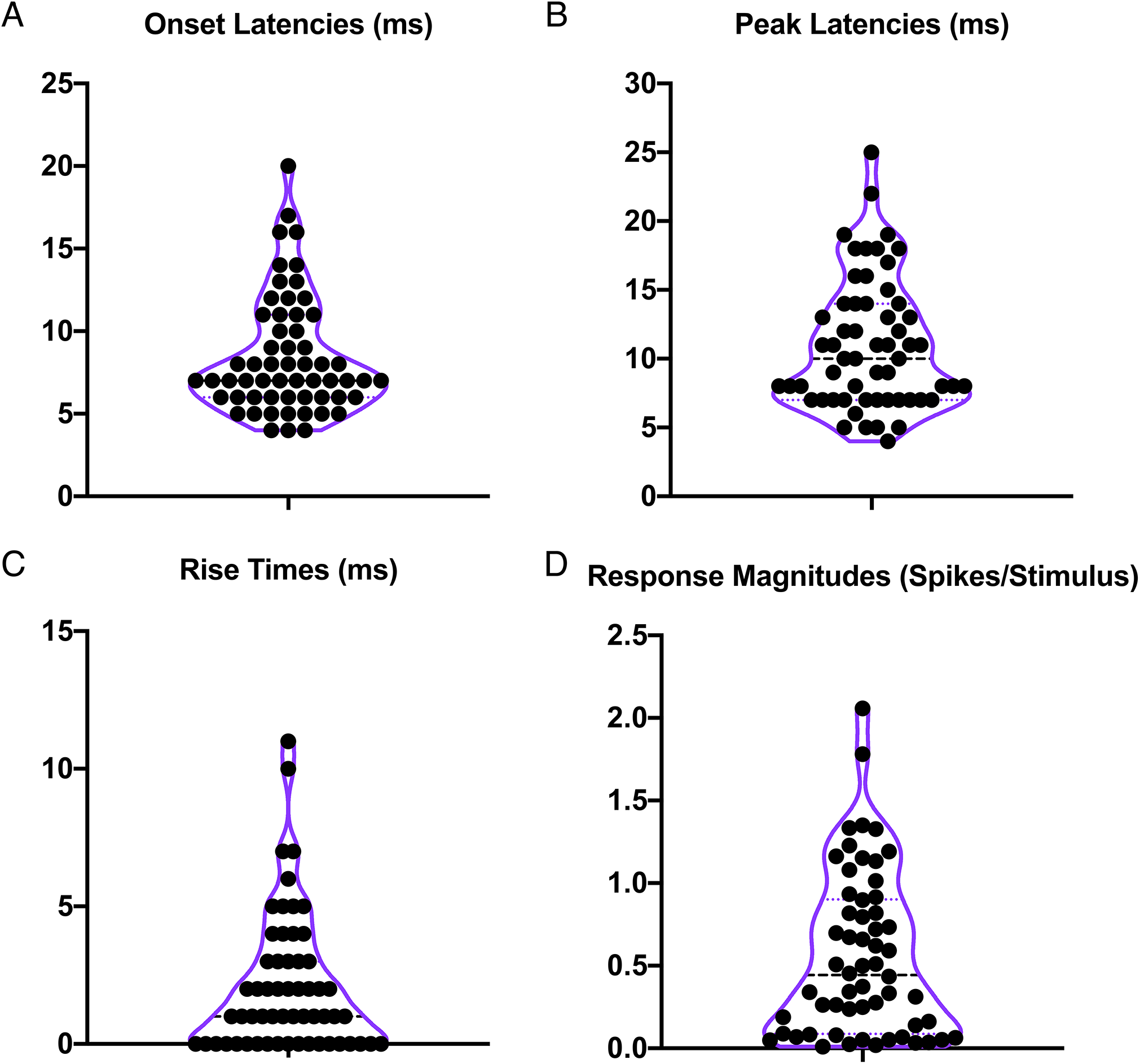
PB neurons respond robustly and with short latency to dura stimulation. (A) The median onset latency is 7ms (95% C.I. 7-8), (B) median peak latency is 10ms (95% C.O. 8 to 12), (C) median rise time is 1ms (95% C.I. 1 - 2). For each panel, n=58 neurons from 22 rats. Each data point represents data from one cell. Dotted horizontal lines depict medians and quartiles. Purple lines show the frequency distribution of the data.

There were no sex differences in peak latency (unpaired t test, t(18) = 0.57, p=0.58), onset latency (unpaired t test, t(18)=0.20, p=0.84), rise time (Mann Whitney U=48.5, p=0.96), or response magnitude (Welch’s t test, t(12.9)=0.32, p=0.75). Because we used rats at a range of ages for the electrophysiology experiments, we tested for age effects by dividing animals into 3 age groups: 2.5 to 3 months (9 rats), 5 to 6 months (5 rats), and 8 to 12 months (7 rats). There were no age differences in peak latency (Kruskal-Wallis statistic = 2.8, p=0.26), onset latency (Kruskal-Wallis statistic = 2.9, p=0.24), rise time (Kruskal-Wallis statistic = 2.4, p=0.31), or response magnitude (Kruskal-Wallis statistic = 1.6, p=0.47).

### 3.2 Dura-responsive PB neurons have diverse receptive fields

We mapped the receptive field of 48 of the 58 recorded neurons. Figure 3 depicts the types of receptive fields we encountered. Most neurons responded only to dura stimulation, and not to noxious or innocuous tactile stimulation anywhere else on the body (28 neurons from both males and females ranging from 3 months to 12 months of age). Four neurons responded to noxious pinch on the hind-limbs bilaterally (from an 8-month old male and a 12-month old female), and 2 responded to pinch on the face bilaterally (from a 3-month old female and a 6-month-old female). One neuron responded to pinch on the hind-limb, fore-limbs, and face bilaterally (from an 8.5-month old male), while two neurons responded to pinch diffusely across the body, with the exception of the tail (from an 8-month old male and a 12-month old female). Eleven neurons responded to pinch across the entire body, including the tail (from two 6-month old males, an 8-month old male, a 3-month old female, a 6-month old female, and an 8-month old female).

**Figure 3:**
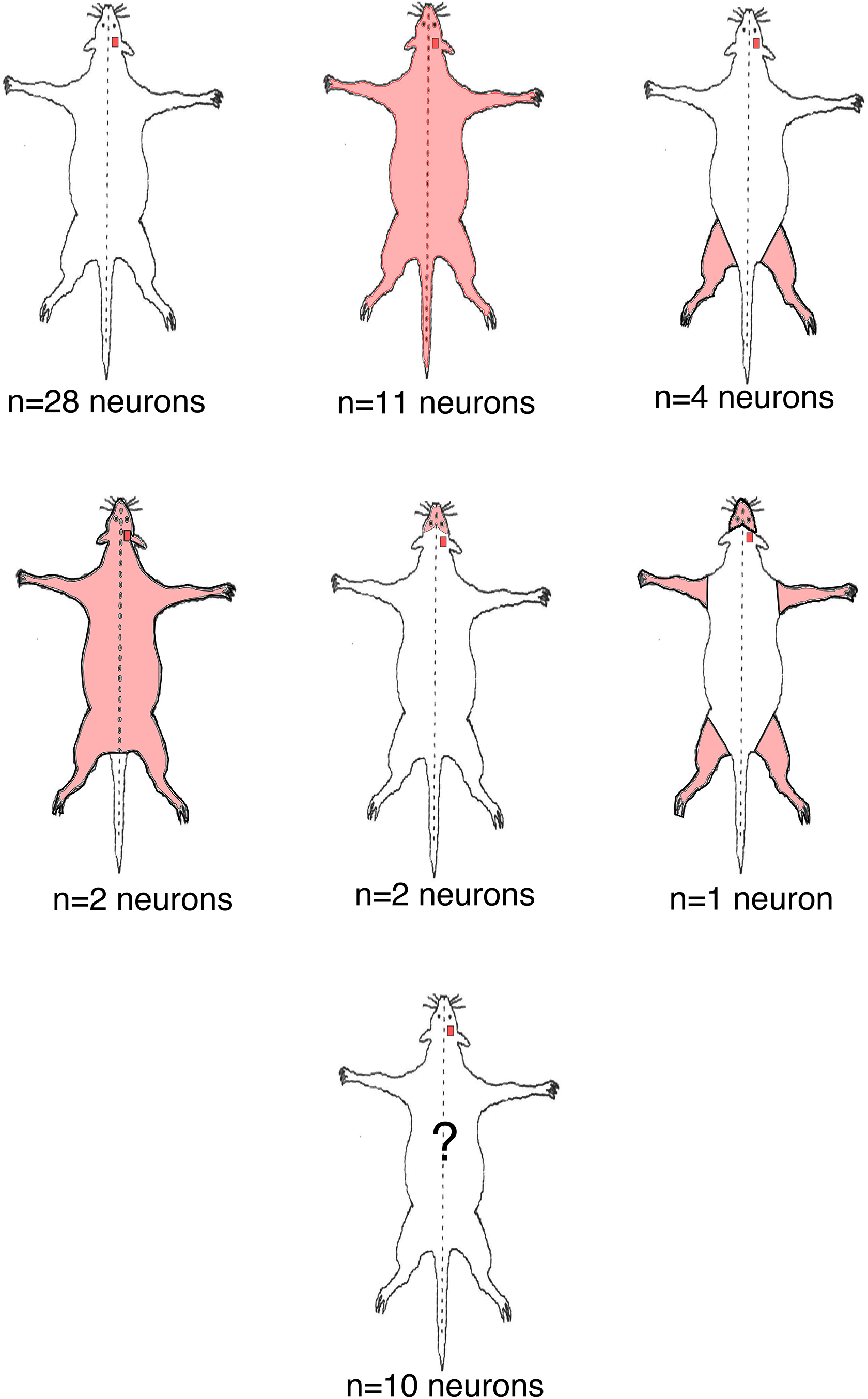
Dura-responsive PB neurons have diverse receptive fields. The small red square represents responses to dura stimulation and pink shading shows the cutaneous receptive field. While most neurons responded only to dura stimulation (28/48), a significant proportion responded both to dura stimulation and to pinch across the entire body (pink shading: 11/48). A small number of neurons responded both to dura stimulation and to pinch on a more restricted receptive field, depicted by the pink shading (n=9 total). We did not obtain receptive field information for 10 of the recorded neurons, denoted by the question mark on the final rat diagram.

### 3.3 Trigeminal ganglion neurons innervating the dura and projecting to PB

After applying fluorescent retrograde tracers to PB and to the dura, we removed and sliced the trigeminal ganglion, examining tissue for evidence of tracer overlap (Fig. 4).

**Figure 4:**
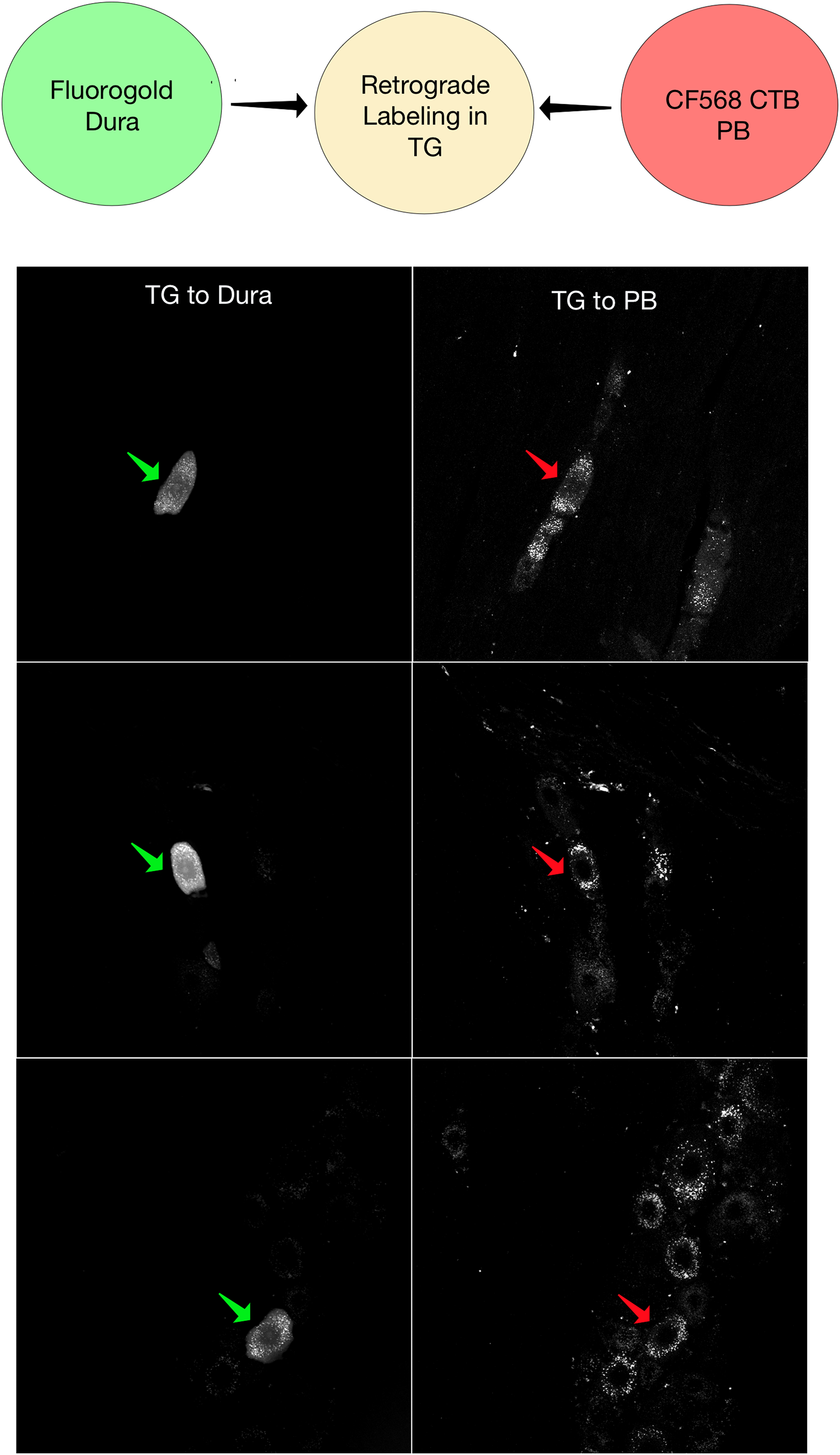
Dura afferents project directly to PB. The diagram depicts the experimental setup. CTB (red arrows) labels PB-projecting TG neurons and Fluorogold (green arrows) labels dura-innervating TG neurons. Examples of dual-labeled neurons are representative images from 3 different animals.

PB injection sites were located primarily in the lateral PB, and resulted in retrogradely-labeled CTB cell bodies in the spinal trigeminal nucleus caudalis and in the nucleus of the solitary tract, nuclei known to project to PB ^36, 37^. In the trigeminal ganglion we identified somata retrogradely labeled with Fluorogold placed on the dura, representing dura-projecting trigeminal afferents (Fig. 4).

We also identified retrogradely-labeled neuronal somata in the trigeminal ganglion, consistent with a direct projection from the trigeminal ganglion to PB (Fig.4). In all animals examined, we identified neurons labeled for both CTB and Fluorogold, representing trigeminal neurons with peripheral axons that innervate the dura, and central axons that project directly to PB (Fig.4). Most Fluorogold labeled somata—representing dura projecting neurons—were interspersed between fiber tracts, presumably belonging to the ophthalmic branch of the trigeminal nerve. Double-labeled somata, representing neurons that innervate both the dura and PB, were located throughout regions containing somata labeled with only Fluorogold, without an obvious somatotopic organization.

## 4. Discussion

### 4.1 Dura-stimulus-responsive PB neurons

We report that PB neurons respond to inputs from the dura. We identified a subset of PB neurons that respond with short latencies to electrical stimulation of the dura mater. These neurons were mostly located in the lateral and external lateral portion of the complex, consistent with evidence of cFos activation in the lateral PB after dura irritation ^38^. The majority of these neurons responded only to dura stimulation, but a considerable number had extensive cutaneous receptive fields. These large receptive fields are consistent with prior reports, including ours, on PB responses ^27,39^. Understanding the receptive fields of these dura-responsive PB neurons is important, because extra-cranial sensitivity —allodynia in particular — is associated with poor treatment response in patients with migraine ^5,6 7^. Further work is needed to clarify how dura responsive PB neurons with wide receptive fields might contribute to extra-cranial pain and allodynia in migraine model conditions.

The short-latency responses of PB neurons are consistent with monosynaptic connections between trigeminal ganglion neurons innervating the dura, and responsive PB neurons. We do recognize that these responses might also reflect a polysynaptic pathway, most likely *via* the spinal trigeminal nucleus caudalis, as there is a known anatomical connection from medullary trigeminal neurons to PB ^37^. However, electrical stimulation of the dura activates medullary trigeminal neurons with an average latency of 11 ms ^40^. Our finding that PB neurons respond to this stimulation with a median onset latency of 7 ms and a median peak latency of 10 ms suggests that these responses are relayed through a direct pathway between the dura and PB.

### 4.2 Anatomical evidence corroborating a direct trigeminal ganglion-to-PB pathway

Several studies suggested a direct connection between the trigeminal ganglion and the parabrachial complex, circumventing the canonical relay site in the spinal trigeminal nucleus ^30–33^. More recently, Rodriguez and colleagues demonstrated a monosynaptic, trigeminal ganglion to PB connection that is implicated in craniofacial pain ^29^. In support of these findings, we demonstrate here retrogradely-labeled trigeminal ganglion neurons after injecting a tracer in PB. These PB-projecting trigeminal ganglion neurons project also to the dura. Thus, these neurons represent a direct pathway between the dura – —a structure implicated in migraine, —and PB –, a key node in chronic pain and aversion.

## 5. Conclusions

Here, we demonstrate short-latency responses to dura stimulation in lateral PB, suggesting a direct pathway between the trigeminal ganglion and PB. Anatomical evidence corroborates this finding, showing trigeminal ganglion neurons that project peripherally to dura and centrally to PB. Given PB’s critical role the critical role of PB’s in maladaptive pain conditions, this pathway is important for further study in the context of migraine pain.

## Abbreviations

PB: Parabrachial complex
TG: trigeminal ganglion
CTB: Cholera Toxin B
FG: FluoroGold = FluoroGold

